# If not a fake, what’s in the lake?

**DOI:** 10.1101/2023.03.01.530639

**Authors:** Floe Foxon

## Abstract

An animal dubbed ‘Champ’ has been sighted by hundreds of eyewitnesses in a large, near-oligotrophic lake in North America. A widely-publicised photograph taken by Mansi purportedly depicting the animal was published to much fanfare. In the present study, sightings were coded and analysed using interrupted time-series models, Pearson correlation coefficients, and descriptive statistics. The number of sightings per year was statistically significantly higher after publication of the Mansi photograph compared to before, which may be evidence of expectant attention, or publicity leading to more lake-goers and therefore more animal sightings. Sightings were consistent in condition (mostly Summer, from Noon to Evening, > 1 witness, and a calm lake surface) which may be interpreted as consistency of when lake-goers visit Champlain, or as evidence of consistent behavioural characteristics of Champ animals. Sightings were highly inconsistent in reported Champ characteristics with widely varying morphology, and most sightings were missing morphological data entirely. More than a quarter of sightings were likened to logs, land mammals, birds, fish, and boats, which are all found in the lake. There were no associations between distance to sighting, estimated length, and estimated height of objects witnessed, which may suggest that eyewitnesses provide inaccurate estimates of these measurements in lake settings. If not a fake, what’s in the lake may be ordinary phenomena mistaken for Champ. Alternatively, Lake Champlain is inhabited by as-yet undiscovered multi-humped, dark-coloured serpents approximately seven meters in length, which locomote in a fast and sinuous fashion, and which enjoy pleasant Summer evenings and crowds. Deciding which explanation best accounts for the data is left as an exercise for the reader.

## Introduction

Lake Champlain is a large, near-oligotrophic lake spanning two US states and one Canadian province, with a maximum depth of 122 meters, and a surface area of over 1,000 square kilometers (Myer, 1979). Not unlike Loch Ness in Scotland, an unknown aquatic animal is alleged by some to inhabit Lake Champlain, variously dubbed ‘The Lake Champlain Monster,’ ‘Champy,’ ‘Champ,’ and the binomen *Beluaaquatica champlainiensis* (a *nomen dubium*, as there is no holotype). Champy has speculatively been identified as outdated reconstructions of such extinct forms as plesiosaurs and archaeocetes (Earley, 1985; Mackal, 1983, Chapter XI).

Perhaps the strongest evidence presented to support the existence of an unknown animal in Lake Champlain is the 1977 ‘Mansi’ photograph, which depicts a dark shape above the lake surface resembling a long-necked marine reptile. Apparently, the picture was not tampered with in any way, but may feature a sand bar upon which a model could have been placed (Frieden, 1984). LeBlond (1982) used the Beaufort scale to estimate the wind speed and wavelength in the photograph, and from these estimated the size of the subject at 4.8 to 17.2 meters. Field experiments have reduced this estimate to 2.1 meters (Radford, 2003) or 1.3 meters (Raynor, 2015). Whatever the size, it has been suggested that the publication of the Mansi photograph may have led to an increase in the number of sightings at the lake due to ‘expectant attention’; the tendency for observers to ‘see’ what they anticipate they will see (Radford and Nickell, 2006, Chapter 2). Thus, one of the aims of the present study is to analyse the impact of the Mansi photograph on Champ sightings statistically.

Throughout the 1980s and early 1990s, the Lake Champlain Phenomena Investigation (LCPI) and associated organisations attempted to collect further evidence with sonar, shoreline and vessel surface surveillance, scuba dives, hydrophones, and a remotely-operated submersible (Deuel and Hall, 1992; Smith, 1984, 1985; Zarzynski, 1982, 1983, 1984b, 1985, 1986b, 1987, 1988, 1989, 1990). Hundreds of eyewitness testimonies were collected as well as a dozen or so ambiguous sonar contacts, but no definitive autoptical evidence was obtained.

In the absense of physical material, purported sightings of unknown animals at Lake Champlain have been studied statistically. In a previously-published analysis, Kojo (1991) reported that Champ sightings most frequently occur just before sunset, in contrast to how most lake visits occur earlier in the day. Kojo interprets this as evidence for the existence of nocturnal unknown animals. However, qualitative descriptions of sightings times such as “late afternoon,” “dusk,” and “about midnight” were excluded, which has the potential to bias the analysis. Thus, a reanalysis including these data is warranted to determine how robust that conclusion is.

The previous analysis reported that the majority of sightings with descriptions of locomotion describe vertical undulation of the body, but did not emphasise how few sightings included any description of locomotion, nor how many sightings were missing data in general. The present study provides exact statistics on the proportion of data missing from sighting reports.

Finally, data on other aspects of sightings such as the diversity of morphological descriptions, the distance to the object, estimates of its size, and whether distance and size are associated in a statistically significant way were not reported in the previous analysis. The present study describes these characteristics.

## Methods

### Data

Most data (224 sightings) were sourced from the definitive Champ book, Zarzynski (1984a, Appendix 4). An additional 58 sightings were sourced from LCPI articles in *Cryptozoology* journal (Zarzynski, 1984b, 1985, 1986b, 1987, 1988, 1989, 1990), and a final 36 sightings not included in the Kojo (1991) analysis were sourced from Champ Quest’s *Cryptozoology* article (Deuel and Hall, 1992), for a total of 318 sightings. Sightings made by the same witness(es) on the same day or shortly thereafter with no details to distinguish between them were coded as the same sighting. Sighting #1, allegedly made by Samuel de Champlain in 1609, was deleted as it has been shown to have resulted from a journalistic error (Radford and Nickell, 2006, Chapter 2). The final analytic sample therefore consisted of *n* = 312 sightings.

Some witnesses reported the month of the sighting, while others reported the season. All months were recoded as seasons (December, January, and February as Winter; March, April, and May as Spring; June, July, and August as Summer; and September, October, and November as Autumn). Similarly, some witnesses reported an exact time of day (e.g., 9 AM) or a range of times (e.g., “2 or 3 PM”), whereas others described the time more broadly, e.g. “afternoon”. Times were recoded as night (22:00–05:59), morning (06:00–11:59, including “morning”), noon/afternoon (12:00–16:59, including “noon”, “late afternoon”, and “early afternoon”), and evening (17:00–21:59, including “evening” and “just before dark”).

Sighting locations were recoded as New York (NY), Vermont (VT), and Quebec (QC). The number of witnesses to a particular sighting was dichotomised: Sightings reportedly seen by a single person were coded as 1 witness, while sightings reportedly witnessed by “several” people, “a party”, “a group”, or any number greater than one were recoded as > 1 witness.

The condition of the lake surface at the time of the sighting was also dichotomised. Conditions such as “like a mirror,” “fine,” “smooth,” “serene,” “still,” “no wind,” and “light breeze” were coded as ‘calm’. Conditions such as “breezy,” “choppy,” “strong wind,” “rough,” “windy,” “ripples,” “wavy,” “waves,” and “rain” were coded as ‘not calm’. Descriptions which detailed only the temperature or sunlight but did not describe the lake surface itself were coded as missing.

Distances to the purported object sighted, lengths of the purported object, and heights above the lake surface of the object were variously reported in yards, feet, miles, inches, and rods. These were all converted to meters and rounded to the nearest 0.1 meters for consistency. “Over 50 ft” was coded as 50 ft, and “less than ¼ mile” was coded as 0.25 miles. Where ranges were given, the midpoint was taken, e.g. “10–12 ft” was recoded as 11 feet. In one case, the size of “a man” was coded as 1.75 meters (the average height of a US male (National Center for Health Statistics, 2021)). In another case, “close enough to give it a whack with the oar” was coded as 3 meters (about the lenth of an oar). “A few” units was coded as 3 units. When the distance changed over the course of the sighting, the value at closest approach was taken. “Close” was considered missing.

The integument of the purported animal was dichotomised as ‘dark’(including “black,” “grey,” “brown,” and “dark green”) and ‘not dark’(including “white” and “light” colours). The general appearance of the body and/or head of the purported animal was categorised as serpentine (including “serpent,” “snake,” and “eel”); reptilian (including “reptile,” “dinosaur” and “alligator”); marine mammal (including “whale,” “manatee,” “seal,” and “dolphin”); land mammal (including “horse,” “moose,” “elephant,” and “dog”); log (including “log,” “tree,” and “pole”); and other (including “periscope,” “stove pipe,” “arm,” “fish,” “bird,” “boat,” “submarine,” and “rubber tube”). The number of humps, loops, coils, or portions of the purported animal was dichotomised as 1 or > 1 (including “several”).

Finally, the speed of locomotion was dichotomised as ‘fast’ (including “great speed,” “railroad speed,” and speeds ≥5 miles per hour) and ‘slow’(including”slow” and speeds <5 miles per hour). Motion was categorised as either ‘sinuous/undulating’ (including “looping” and “up and down”), or ‘rolling’.

### Analyses

To investigate the impact of the highly-publicised Mansi sighting and photograph on purported sightings of Champ at Lake Champlain, interrupted time-series analyses were undertaken. Linear fixed-effects regression models were implemented, which regressed the cumulative number of Champ sightings on time (year) to provide a statistical estimate of the average number of sightings per year with 95% confidence intervals. This analysis was repeated for the years preceeding the publication of the Mansi photograph, and for the years following publication. By comparing the confidence limits on the estimates for each time-series, it is possible to determine whether the Mansi publication was associated with a statistically significant change in the number of sightings. Note that while the Mansi sighting purportedly took place on July 5, 1977, the photograph and sighting details were not published publicly until June 30, 1981 (Radford and Nickell, 2006, Chapter 2). Therefore, the latter date was used as the cut-off for the time-series analyses. The Mansi sighting was necessarily removed for these analyses.

Descriptive statistics on the distribution of sightings by season, time, state, number of witnesses, lake surface condition, distance, length, height above lake surface, integument, appearance, speed, and motion are tabulated for the re-coded data, with the proportion of data missing also reported. For distance, length, and height variables, the median and interquartile range, as well as the mean and standard deviation are reported. Means and standard deviations were calculated excluding outliers. Outliers were identified using the bestfitting probability density functions for each of these variables as in Taskesen (2023).

To investigate the possible association between distance, length, and height, Pearson correlation coefficients were calculated.

All analyses were performed in Python 3.8.16 with the packages Numpy 1.21.5, Pandas 1.5.2, Distfit 1.6.6, Scipy 1.7.3, Statsmodels 0.13.5, and Matplotlib 3.6.2. All code and data are available in the online Supplementary Information (Foxon, 2023).

## Results

Figure 1 shows the cumulative number of sightings of unknown animals at Lake Champlain across the period of time ±10 years around the 1977 Mansi photograph (published in 1981). As detailed in the figure, numbers of sightings per year were over twice as great in the periods 2, 5, and 10 years *after* the publication of the Mansi photograph in 1981, compared to the periods 2, 5, and 10 years *before* publication. The increase in number of sightings per year after the Mansi photograph diminished around 1987/8 when sightings per year returned to pre-Mansi levels.

**Figure 1.**
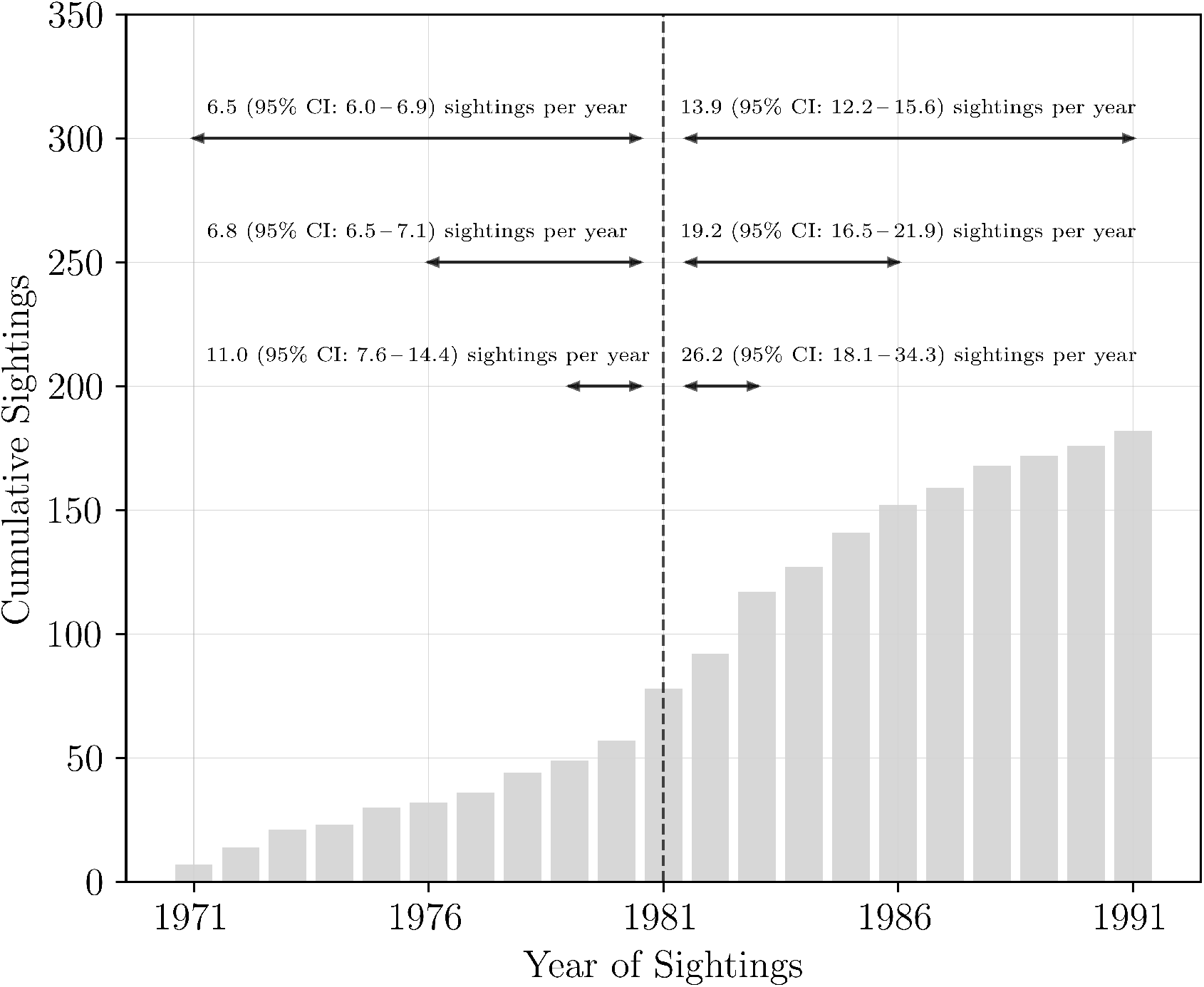
Cumulative number of sightings of unknown animals at Lake Champlain ±10 years around the publication of the Mansi photograph on June 30,1981. The vertical dashed line indicates the year in which the Mansi photograph was published.

Table 1 shows the distribution of all *n* = 312 sightings by sighting conditions and purported characteristics. The conditions under which sightings were made were largely consistent; the majority of sightings occured in Summer (70.1%) under calm lake surface conditions (83.9%), between Noon and Evening (73.9%), and were made by more than one witness (72.9%). However, descriptions of the characteristics of the animal observed are much less consistent. Just 27% of witnesses described the appearance of the animal they had purportedly seen. Among those reports with appearance descriptions, the largest category (serpentine) consistent of just 36.5% of sightings, with the next largest specific categories being reptilian (15.3%) and marine mammal (11.8%). Over 68% of sightings were missing data on one or more of appearance, motion, speed, height above the lake surface, and the number of humps, loops, coils, or portions seen. The characteristic with the most agreement was integument;98.2% of sightings with data for integument described the animal as dark, which is unsurprising in a lake setting.

**Table 1.**
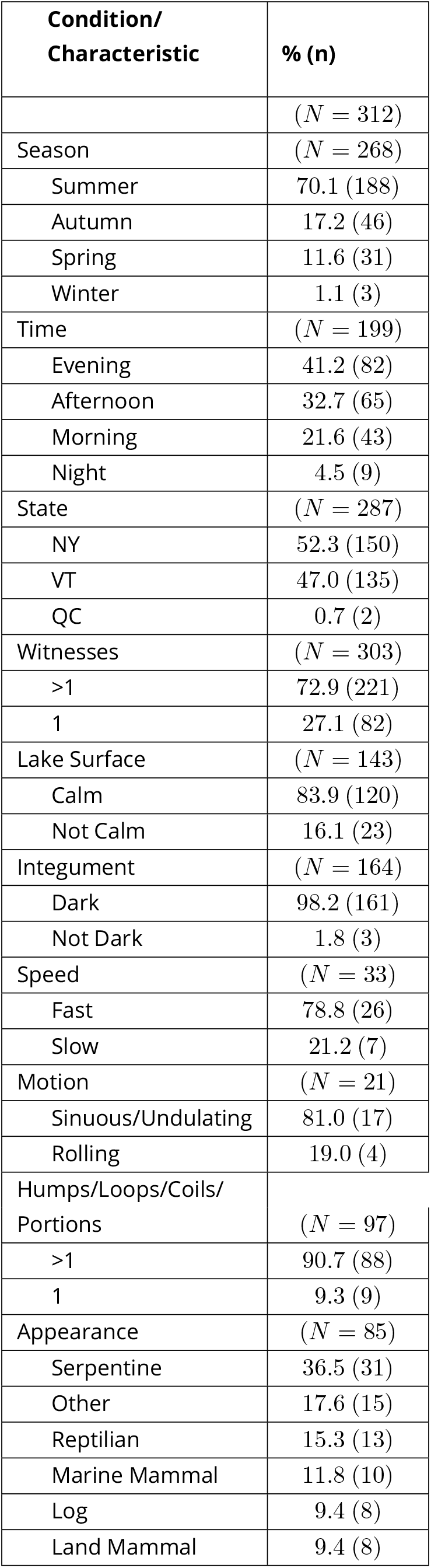
Conditions under which purported sightings of unknown animals at Lake Champlain were made, and reported characteristics of those animals.

The average reported distance between observer and object was 247.2 (standard deviation: 6.5) meters, and the median distance and interquartile range was 91.4 (240.8) meters. For length the average was 6.6 (3.0) meters and the median was 6.9 (4.5), and for height the average was 1.1 (0.7) meters and the median was 0.9 (0.9) meters. The only statistically significant correlation between the continuous variables was between the reported length and height of the animal when outliers were included (*r* = 0.623, *p* < 0.001), but this association disappeared when outliers were excluded (*r* = 0.162, *p* = 0.402). Correlations between distance and length (*r* = –0.004, *p* = 0.969 incl. outliers;*r* = 0.093, *p* = 0.402 excl. outliers) and between distance and height (*r* = 0.177, *p* = 0.245 incl. outliers;*r* = 0.246, *p* = 0.107 excl. outliers) were near zero and statistically non-significant.

## Discussion

This study presents the most thorough statistical analysis of purported sightings of unknown animals at Lake Champlain to date. It is shown that while sightings mostly occur under similar conditions (Summer, Noon to Evening, > 1 witness, and calm lake surface), eyewitness descriptions of these alleged animals are few and disparate. Most sightings are missing data pertaining to morphological description, and those with data describe very different-looking objects. Thus, Champ sightings are not nearly as consistent as claimed by some (Radford and Nickell, 2006, Chapter 2). That a large proportion (27.4%) of sightings were described as appearing like logs, land mammals, birds, fish, and boats is logical because all of these are present at Lake Champlain, for example in 1987 the LCPI observed deer swimming across the lake. Unknown animals are therefore unnecessary to explain many sightings. Sightings that occured in late evening with low visibility are perhaps explained by observers being more likely to confuse ordinary phenomena with Champy in low light conditions.

The finding that there are no associations between distance, length, and height may suggest that some eyewitnesses over- or underestimate these factors, or are witnessing different events occuring in the lake.

The finding that the rate of sightings increased significantly after the publication of the Mansi photograph may be interpreted by a cryptozoologist as evidence of publicity driving more cryptid-seekers to Lake Champlain, and therefore more sightings being made (since the number of sightings is a function of the number of visitors). A skeptic may interpret the same finding as evidence of expectant attention; after seeing the Mansi photograph, lake-goers may be more likely to think they have seen Champ because that is what they are anticipating.

Similarly, a cryptozoologist may interpret the consistency of sighting conditions as evidence of consistent behavioural characteristics among Champ animals, while a skeptic may interpret this consistency as evidence of regularity of wind/wave effects (Bauer, 1992), sampling error or other bias, and/or characteristics not of the Champ animals but of the observers (Mackal, 1992). Indeed, in the case of the cryptozoologically-related coelacanth, catch data describes more about the habits of Comoran fisherman than about the fish (Thomson, 1992, Chapter 6). For example, Kojo (1991) suggested that most Champ sightings occuring in Summer is evidence of seasonal heterothermy, but this may also be explained by seasonal tourism in and around Lake Champlain.

In conclusion, if not a fake, what’s in the lake may be ordinary phenomena innocently mistaken for unknown animals, in part driven by expectant attention due to the publicity of the Mansi photograph. Alternatively, Lake Champlain is inhabited by multi-humped, dark-coloured serpents approximately seven meters in length, which locomote in a fast and sinuous fashion, and which prefer pleasant Summer afternoons and evenings, as well as appearing before crowds. Deciding which explanation best accounts for the data is left as an exercise for the reader. Regardless, Champ research and advocacy has apparently benefitted the local people and animals; in 1986, the Vermont Senate followed the New York Senate in adopting the “Champ Resolution,” calling for the protection of possible unknown animals in the lake, which likely benefited the known lake fauna. Systematic searches at Lake Champlain have also enabled the discovery of many archaeological artefacts (Zarzynski, 1986a) such as the shipwreck of the tugboat William McAllister, which was discovered off Schuyler Island during a six-day search of the lake bed designated ‘Project Champ Carcass’ (Zarzynski, 1987).

## Data, script, code, and supplementary information availability

Data and code are available online: https://doi.org/10.17605/OSF.IO/C34WY

